# Sequence reliance of a *Drosophila* context-dependent transcription factor

**DOI:** 10.1101/2023.12.07.570650

**Authors:** Lauren J. Hodkinson, Julia Gross, Casey A. Schmidt, Pamela P. Diaz-Saldana, Tsutomo Aoki, Leila E. Rieder

## Abstract

Despite binding similar *cis* elements in multiple locations, a single transcription factor often performs context-dependent functions at different loci. How factors integrate *cis* sequence and genomic context is still poorly understood and has implications for off-target effects in genetic engineering. The *Drosophila* context-dependent transcription factor CLAMP targets similar GA-rich *cis* elements on the X-chromosome and at the histone gene locus but recruits very different, loci-specific factors. We discover that CLAMP leverages information from both *cis* element and local sequence to perform context-specific functions. Our observations imply the importance of other cues, including protein-protein interactions and the presence of additional cofactors.

## Introduction

Coordinated gene expression is a crucial but difficult task in the crowded nucleus. To accomplish this feat, transcription factors (TFs) must first traverse the nucleus to find their corresponding *cis* elements. Furthermore, once factors have identified their DNA-binding sites, they can then impact gene expression on highly constrained temporal and spatial levels. When gene expression programs are misregulated or interrupted by mutations in regulatory elements, it can have catastrophic impacts, causing a variety of disease outcomes including cancer, autoimmunity, and neurological disorders (Lee and Young 2013). To further complicate the hurdle of widespread gene regulation, some TFs are “context dependent”; they bind similar *cis* elements across the genome but retain the ability to perform distinct functions at these different loci (Fry and Farnham 1999). Currently, we still do not fully understand how context-dependent TFs integrate locational information with cues they may receive from cofactors, 3D architecture, and other signals to perform their diverse functions.

CLAMP (Chromatin Linked Adaptor for MSL Proteins) is *Diptera*-specific C2H2 zinc-finger TF that plays a genome-wide role as a pioneer factor (Duan et al. 2021) and targets GA-rich *cis* elements to regulate global gene expression through both chromatin accessibility changes and polymerase pausing (Urban et al. 2017b, 2017a). CLAMP is designated as a context-dependent transcription factor that is critical for two major coordinated gene expression events: histone gene expression and upregulation of the male X chromosome for dosage compensation (Soruco and Larschan 2014). CLAMP is enriched on the male X-chromosome where it binds to GA-rich regions called MREs (MSL recognition elements) (Alekseyenko et al. 2008) and recruits the Male Specific Lethal complex (MSLc) to accomplish dosage compensation in male flies (Soruco and Larschan 2014). At the histone gene locus, CLAMP binds a long GA-repeat *cis* element in the promoter of *H3* and *H4* (*H3/H4*p) and fosters recruitment of histone gene locus body (HLB) specific factors including Mxc (Multi Sex combs, Mxc; the *Drosophila* ortholog of the human NPAT; (Terzo et al. 2015)) (Salzler et al. 2013; Rieder et al. 2017). CLAMP targets both the histone genes and the X chromosome prior to locus-specific factors such as Mxc and MSLc, neither of which have strong DNA-binding capability (Villa et al. 2016; Terzo et al. 2015). These observations show that, despite binding similar *cis* elements on the X-chromosome and at the histone gene locus, CLAMP recruits different factors to each location ensuring proper group composition at each genomic location. It is unclear how early transcription factors such as CLAMP integrate information from GA-rich *cis* elements with other genomic contextual information to perform their locus-specific functions.

*Drosophila* dosage compensation provides some clues as to how TFs like CLAMP integrate sequence and context information. For example, moving X-linked chromosome entry sites (CES; up to ∼1500bp which contain one or more GA-rich MREs (Alekseyenko et al. 2008)) to autosomal locations leads to ectopic MSLc recruitment, spreading of the complex into surrounding chromatin, and transcriptional regulation of nearby genes (Gorchakov et al. 2009). A similar phenomenon occurs when the ∼300bp *H3/H4*p, which includes a GA-repeat ranging from 16-35 bps (Bongartz and Schloissnig 2018) targeted by CLAMP (Salzler et al. 2013; Rieder et al. 2017). When this segment is placed outside the endogenous histone gene locus on chromosome 2L, HLB-specific factors are recruited to the transgenes resulting in transcription of the adjacent sequences (Salzler et al. 2013). These important experiments show that separation of local contexts (CES, up to ∼1500bp; *H3/H3*p, ∼300bp) from the larger locus (X-chromosome, histone gene locus on chromosome 2L) still allows for retention of local context function. This retention of function suggests that the larger chromosomal or locus context of these elements is not required for their function in recruiting the factors necessary for coordinated gene expression. Since both regions carry elements that recruit the CLAMP protein (Alekseyenko et al. 2008; Rieder et al. 2017), we hypothesized that the GA *cis* elements themselves are interchangeable and that the flanking local context provides the cues required for CLAMP context-specific function and unique factor recruitment.

Neither the *Drosophila* X-chromosome nor the endogenous histone gene locus are tractable study systems in which to test this hypothesis. MSLc coats the entire chromosome, and it is not practical to manipulate each GA-rich MRE in all 150 CES (Alekseyenko et al. 2008). Altering a critical number of CES, besides being nearly impossible to execute, would likely cause incomplete dosage compensation and male-specific lethality, while altering just a few CES is unlikely to significantly affect MSL recruitment to the whole chromosome due to complex spreading (Kelley et al. 1999; Kageyama et al. 2001; Gorchakov et al. 2009). Similarly, the endogenous histone gene locus is organized in a series of ∼100 tandemly repeated 5 Kb arrays, in which each array contains the five canonical histone genes (*H3, H4, H2A, H2B*, and *H1*).

The repetitive nature of the histone locus renders it intractable for genetic manipulation as it harbors CLAMP-binding GA-repeats in all ∼100 *H3/H4*p. However, the *Drosophila* transgenic histone gene array provides an excellent genetic system in which to test our hypothesis. Transgenes carrying 1-12 histone gene arrays have been established that allow for genetic manipulation and recapitulate histone locus functionality (Salzler et al. 2013; McKay et al. 2015; Meers et al. 2018). A single copy histone gene array, while not able to rescue an endogenous histone locus deletion background, has been shown to successfully recruit HLB-specific factors and drive histone gene expression (Koreski et al. 2020). This becomes a powerful system to perturb the CLAMP-recruiting *cis* elements within the array without editing all ∼100 arrays at the endogenous locus.

Leveraging the transgenic system, we confirm that *H3/H4*p GA-repeat CLAMP binding sites are required for Mxc recruitment to the transgene (Rieder et al. 2017). We further demonstrate that transgenes in which we replace the *H3/H4*p GA-repeats with X-linked GA-rich MREs, which are bound by CLAMP in vitro and in their native locations, fail to recruit Mxc despite the larger contextual information of the histone gene array. Finally, we demonstrate that adding back additional X-linked sequence to the transgenic histone gene array results in MSLc recruitment in males. We observe sex-specific differences that suggest a competition between CLAMP-associated factors in males, but not in females. Overall, our observations indicate that *cis* element sequence alone is enough to impact context-dependent TF functions.

## Results and Discussion

### CLAMP targets GA-rich *cis* elements at different loci

CLAMP binds genome-wide and is enriched on the X-chromosome as well as on chromosome 2L at the endogenous histone locus (**Figure 1A**). We mapped available CLAMP ChIP-seq data from 2-4 hr mixed embryos (Duan et al. 2021) and confirmed that CLAMP targets the GA-repeats in the *H3/H4*p (**Figure 1B**). Mxc is specific to the histone locus by immunofluorescence (White et al. 2011) and the mammalian homolog NPAT also only targets histone promoters in cultured human cells (Kaya-Okur et al. 2019). We confirmed that Mxc is specific to the histone genes by performing ChIP-seq from embryo samples and by performing sexed polytene chromosome immunofluorescence using antibodies against Mxc, CLAMP, and MSL3. Mxc targets only the histone locus on chromosome 2L (**Figure 1A, D**) and is broadly localized over histone gene promoters, overlapping with CLAMP signal at the *H3/H4p* (**Figure 1B**). We also mapped existing MSL3 (Male Specific Lethal 3; a component of MSLc) ChIP-seq data from embryos (Rieder et al. 2019) and confirmed that MSLc is enriched on the X-chromosome (**Figure 1A, D**). As expected, MSLc is not enriched at the autosomal histone locus (**Figures 1A-B, D**). Both CLAMP and MSL are enriched at CESs, including CES5C2 and the *roX2* (*RNA on X 2*; a lncRNA component of MSLc) CES (**Figure 1C)**. Although CLAMP is present at both X-linked CES and the GA-repeats within the context of the histone gene array, MSLc and Mxc are locus-specific.

**Figure 1:**
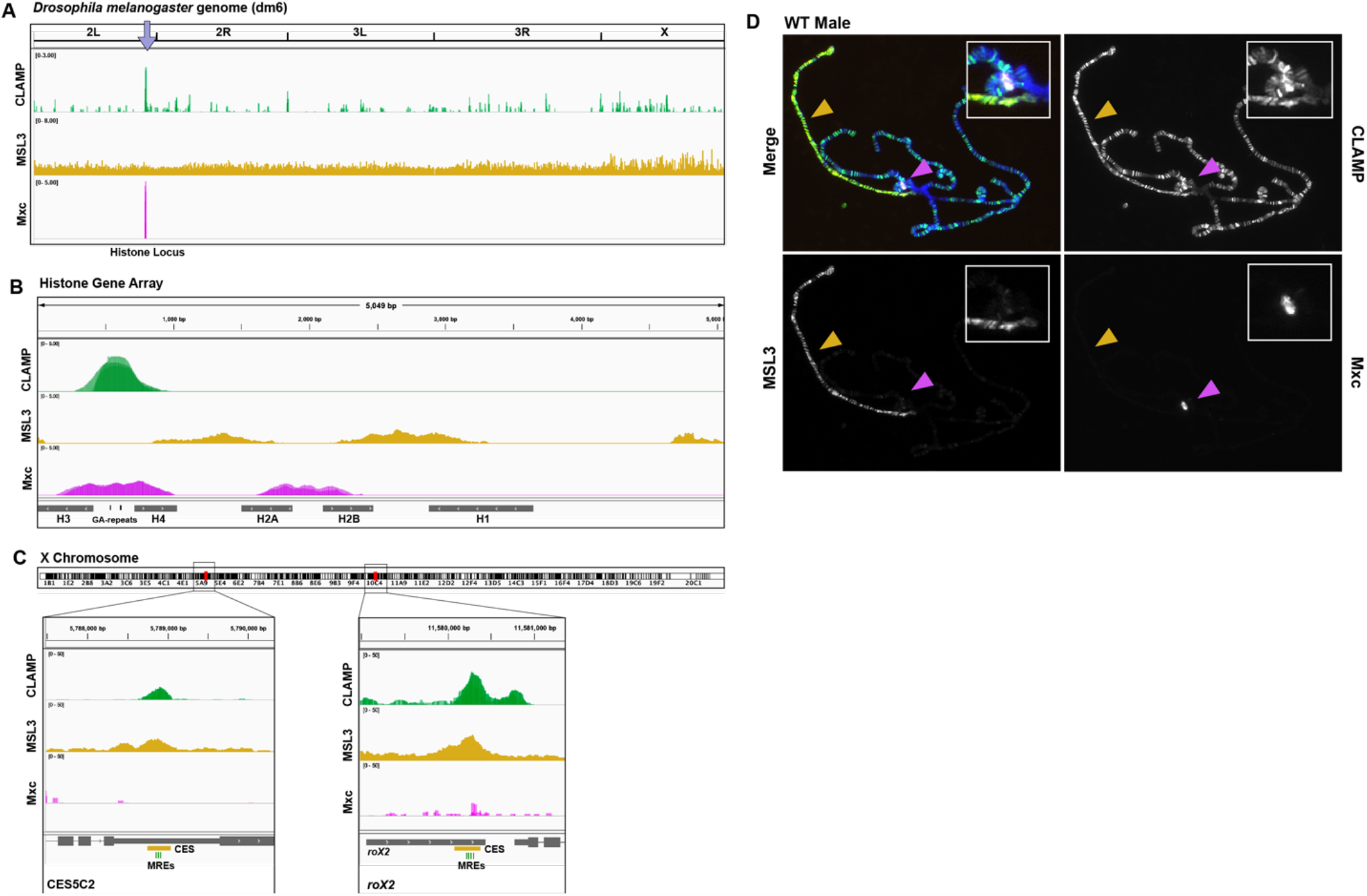
CLAMP binds locations genome-wide while Mxc and MSL3 bind distinct genomic regions. **(A)** We mapped ChIP-seq data for CLAMP (green) from 2-4hr mixed sex embryos (Duan *et al*. 2021, GSE152598, three overlaid replicates normalized to respective inputs), MSL3 (yellow) from 2-4hr mixed embryos ((Rieder et al. 2019), GSE133637, normalized to the input), and Mxc (magenta) from 2-4 hr female staged embryos (three overlaid replicates normalized to respective inputs) to the dm6 (*Drosophila melanogaster)* genome. The purple arrow indicates the location of the endogenous histone gene locus. **(B)** CLAMP, MSL3, and Mxc ChIP-seq data mapped to a custom single histone gene array. The dark green bars between the *H3* and *H4* coding sequences mark the CLAMP binding GA-repeats. **(C)** ChIP peaks at two X-chromosome locations, CES5C2 and the *roX2* gene. The yellow bars mark the known chromosome entry sites (CES) and the light green bars mark the GA-rich MSL recognition elements (MREs) where CLAMP and MSLc colocalize. **(D)** We performed immunofluorescence staining of wildtype chromosomal male third instar larval polytene chromosomes for CLAMP (green), MSL3 (yellow) and Mxc (magenta). DNA is stained with DAPI (blue). CLAMP and Mxc colocalize at the endogenous histone locus (magenta arrow, solid outlined box) and CLAMP and MSL3 colocalize on the X-chromosome (yellow arrow). Mxc and MSL3 do not colocalize.

### CLAMP requires GA-rich sequences to bind *in vitro* and *in vivo*

CLAMP is a zinc-finger protein that directly interacts with DNA sequence (Soruco and Larschan 2014). Recombinant full-length CLAMP binds to GA-repeat carrying DNA probes *in vitro* (Duan et al. 2021). We therefore investigated the ability of recombinant CLAMP to interact with different *cis* element and whether the GA-repeats were critical for CLAMP binding *in vitro*. We first designed two biotin-labeled DNA probes based on the sequence from the endogenous *H3/H4*p. The “WT” probe includes the endogenous *H3/H4*p with intact CLAMP-recruiting GA-repeat. The “GA delete” probe includes the same *H3/H4*p sequence except the GA-repeats are removed (**Figure 2A**). CLAMP is maternally deposited in the early embryo and is a known member of the Late Boundary Complex which shifts CES probes *in vitro* (Kuzu et al. 2016; Kaye et al. 2017). We therefore also performed radiolabeled EMSAs using embryo extracts, which should include both maternally deposited CLAMP as well as the CLAMP-containing LBC, and our probes. To confirm the embryo extract included CLAMP, we performed shifts with extract, probe, and CLAMP antibody and showed that CLAMP antibody super-shifted with the WT WT probe (**Supplemental Figure 1)**. Only the WT probe containing the GA-repeats is shifted with embryo extract; the GA delete probe did not shift (**Figure 4B**). We repeated these EMSAs with recombinant full-length CLAMP protein (Duan et al. 2021) with the same result (**Figure 2B**). These observations confirm that the endogenous *H3/H4*p GA-repeats are critical for CLAMP binding *in vitro*.

**Figure 2:**
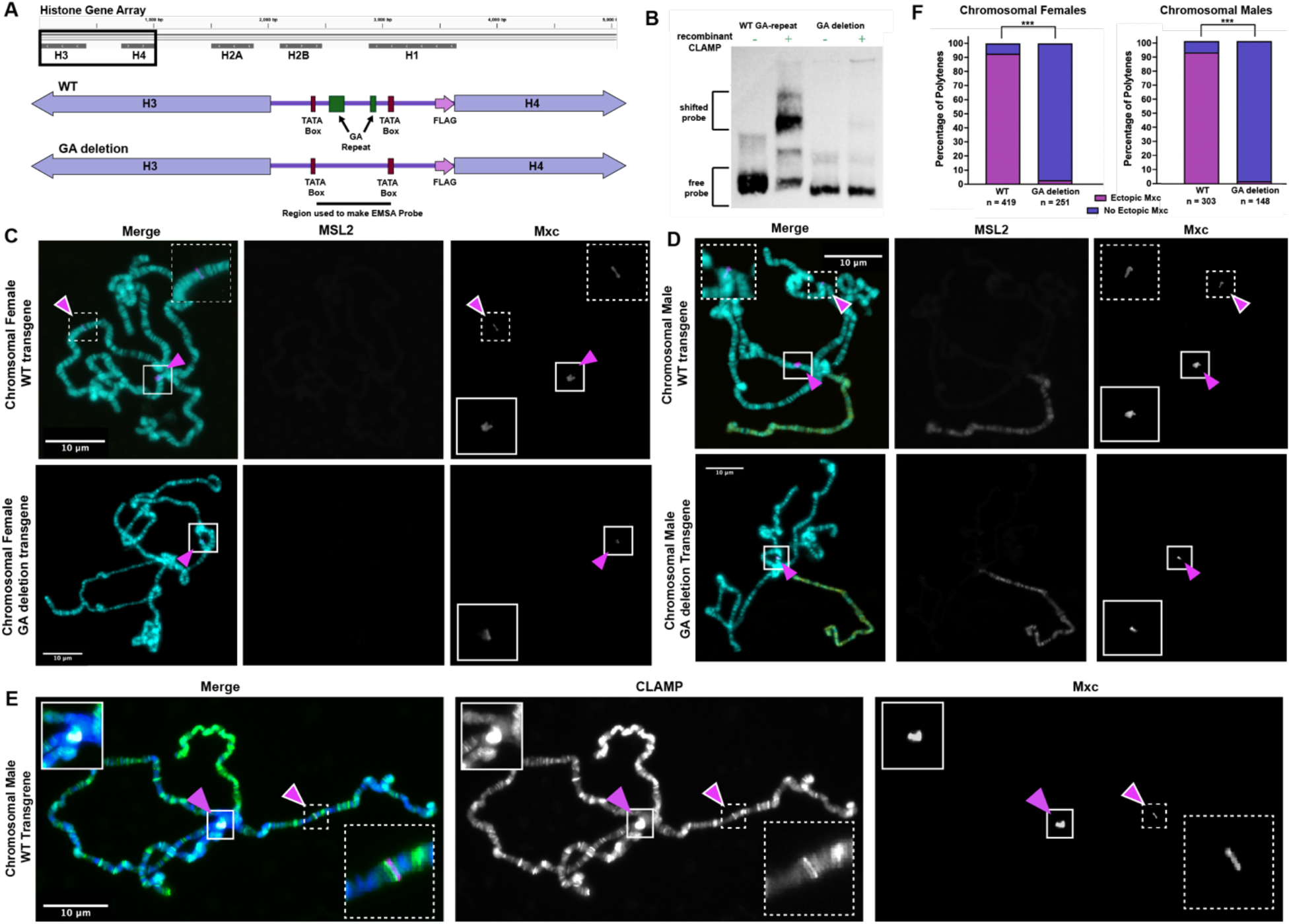
GA-repeats are required for CLAMP binding and function at the transgenic histone gene array. **(A)** We engineered two transgenes carrying a single histone gene array: the “WT” transgene which resembles the endogenous histone arrays and the “GA deletion” transgene in which we deleted the GA-repeats (dark green, labeled bars). **(B)** We performed EMSA (gel shift) assays with recombinant CLAMP and biotinylated probes of the *H3/H4p* sequences from both histone array transgenes (WT: 226 bp, GA deletion: 198 bp). Recombinant CLAMP shifts EMSA probes only when the GA-repeats are present. **(C)** We performed immunofluorescence staining of third instar larval polytene chromosomes in chromosomal females for Mxc (magenta). A chromosomal female carrying the WT transgene (top) shows ectopic Mxc (white outlined magenta arrow, dotted outlined box) and while a chromosomal female carrying the GA deletion transgene (bottom) shows no ectopic Mxc staining. **(D)** A chromosomal male carrying the WT transgene (top) shows ectopic Mxc while a chromosomal male carrying the GA deletion transgene (bottom) shows no ectopic Mxc staining. Both sexes show Mxc localizing to the endogenous histone locus, which is used as an internal staining control (magenta arrow, solid outlined box). **(E)** We also performed immunofluorescence in chromosomal male and female animals for CLAMP (green) and Mxc (magenta). A chromosomal male shows colocalization of CLAMP (green) and Mxc (magenta) at the endogenous histone locus (magenta arrow, solid outlined box) and at the WT transgene (white outlined magenta arrow, dotted outlined box). Both chromosomal females and chromosomal males show Mxc localizing to the endogenous histone locus, which is used as an internal staining control (magenta arrow, solid outlined box). **(F)** Quantification of ectopic Mxc from polytene scoring shows a significant difference between the percentage of chromosome spreads that have ectopic Mxc between the WT and GA deletion transgenes in both chromosomal males and chromosomal females. n values reflect number of polytenes scored for each respective genotype. *** Chi-squared text, p < 0.001

*In vivo*, GA-repeat *cis* elements are clearly not sufficient for CLAMP and Mxc recruitment: GA-repeats exist throughout the genome and long repeats are enriched on the X-chromosome (Kuzu et al. 2016) yet Mxc is solely recruited to the histone locus (**Figure 1**) (Rieder et al. 2017, 2019). We therefore sought to translate our *in vitro* observations *in vivo*. A transgene carrying twelve wild-type histone gene arrays recruits HLB components, including CLAMP and Mxc, but transgenic arrays lacking the GA-repeats in the *H3/H4*p fail to attract HLB factors when the endogenous locus is present (Rieder et al. 2017; Koreski et al. 2020). To validate these findings and to confirm these observations using a transgene carrying only a single histone gene array, we created two transgenic lines. The “WT” line carries a transgene with a single wild-type copy of the histone gene array, while the “GA deletion” line carries the same transgene lacking the GA-repeat sequences in the *H3/H4*p (**Figure 2A**). We then performed sexed polytene chromosome immunofluorescence using an antibody against Mxc and scored for ectopic Mxc.

We observed that both CLAMP and Mxc are recruited to the WT transgene in both chromosomal female and chromosomal male larvae (**Figure 2C**,**D**,**E**). The endogenous histone locus is visible near the chromocenter and serves as an internal staining control. Approximately 80% of polytenes from female larvae containing the WT transgene and 85% of polytenes from male larvae exhibited ectopic Mxc recruitment (**Figure 2C**,**D)**. In contrast, we rarely observed ectopic Mxc recruitment in larvae carrying the GA delete transgene (**Figure 2C**,**D**,**E**) and observed a significant difference between the polytene scoring of WT larva and GA-deletion larva (**Figure 2F)**. Our *in vivo* observations are therefore in agreement with our *in vitro* EMSA results and establish the transgenic system as a manipulable *in vivo* assay.

Because CLAMP is an integral transcription factor responsible for regulating the expression of essential genes such as the histone genes, it’s likely that there are “backup” mechanisms for ensuring CLAMP can locate regions where GA-rich *cis* elements reside and perform its function. Recent work demonstrated that CLAMP is recruited to a transgene carrying twelve histone gene arrays in which the *H3/H4*p is replaced with the *H2A/H2B* promoter, which does not contain GA-repeats. But, the phenomenon where CLAMP and HLB factors are recruited to the transgene only occurs in the background of a endogenous ∼100 copy histone locus deletion (Koreski et al. 2020). CLAMP targets the region by immunofluorescence, but does not interact with any sequence by ChIP-seq. This is still surprising, since our observations show that simply deleting the GA-repeats from the *H3/H4*p rendered CLAMP non-functional in the context of the single histone array transgene (**Figure 2C**,**D**). However, all of our observations are by necessity in the background of the endogenous histone locus, as the single histone gene array itself does not support viability (Günesdogan et al. 2010; McKay et al. 2015).

### The GA-repeats must reside in the *H3/H4*p for Mxc recruitment in vivo

We next sought to determine if simply the local promoter context (∼300 bp) affects CLAMP recruitment to target elements. We hypothesized that the GA-repeats could be moved anywhere within the transgenic histone gene array and still attract CLAMP along with other histone locus specific factors, such as Mxc, because the larger context of the array is retained. We engineered two transgenes in which we deleted the GA-repeats from the *H3/H4*p and moved them to one of two locations within the transgenic array: either within the *H2A/H2B* promoter (“*H2A/H2B*”) or within the intergenic region between the *H1* and the *H2B* coding sequences (“*H2B/H1*”) (**Figure 3A**). In both cases, we attempted to avoid disrupting any known or predicted *cis* elements (Crayton et al. 2004). We then performed fluorescent staining on polytene chromosomes from transgenic lines with an antibody against Mxc to assess how the different GA-repeat locations impacted Mxc recruitment compared to controls (**Figure 3B**,**C)**. We rarely observed ectopic Mxc in transgenic lines in which the GA-repeats reside in the *H2A/H2B* promoter or the region between *H1 and H2B* (**Figure 3C**,**D**). Our data showed a significant difference between the percentage of chromosome spreads with ectopic Mxc for both transgenes when compared to our WT transgene and the data more closely resembled that of the GA deletion transgenes in which we completely removed the GA-repeats (**Figure 3D)**. Our observations show that neither the *H2A/H2B* or *H2B/H1* transgenes led to ectopic Mxc recruitment, indicating the importance of local *H3/H4*p context for Mxc recruitment.

**Figure 3:**
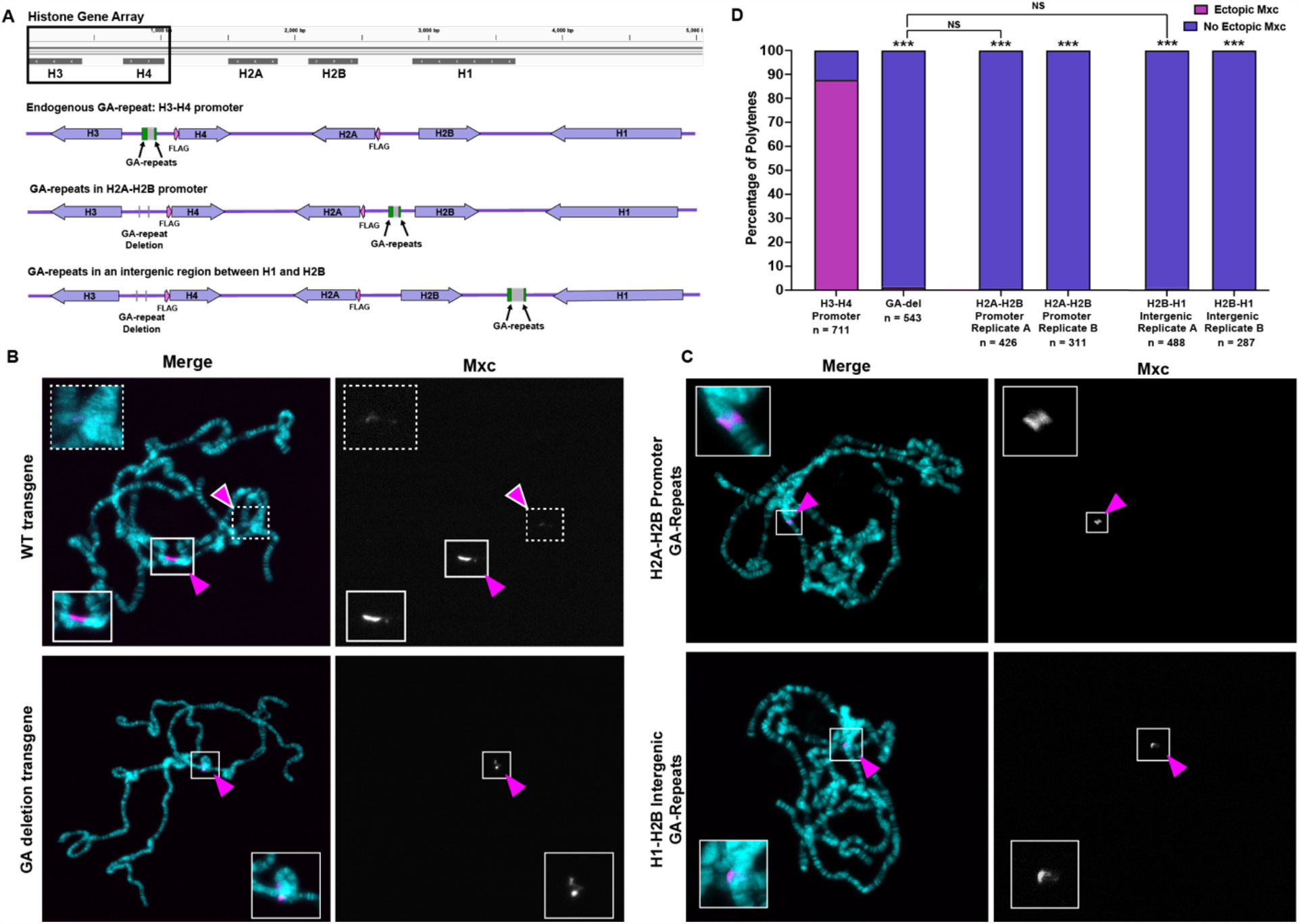
GA-repeat*s* must reside in the *H3/H4*p for proper CLAMP function at the transgenic histone gene array. **(A)** We engineered two histone gene array transgenes *in which* we moved the GA-repeats (green bars) to different regions along the array; one where we placed the GA-repeats in the *H2A/H2B* promoter and one where we placed the GA-repeats in the intergenic region between *H1* and *H2B*. **(B)** We performed immunofluorescence staining *of* third instar larval polytene chromosomes for Mxc (magenta). DNA is stained with DAPI (cyan). Animals carrying the WT transgene (top) show ectopic concentration of Mxc (white outlined magenta arrow, dotted outlined box) while animals carrying the GA deletion transgene (bottom) show no ectopic Mxc staining. **(C)** Animals carrying the transgenes in which the GA-repeats are moved to the *H2A/H2B* promoter (top) or intergenic region between *H1* and *H2B* (bottom) do not show ectopic Mxc. All animals show Mxc localizing to the endogenous histone locus, which is used as an internal staining control (magenta arrow, solid outlined box). **(D)** Quantification of ectopic Mxc from polytene scoring shows a significant difference in the percentage of chromosome spreads that have ectopic Mxc between WT and the transgenes in which the GA-repeats are moved. Significance above the bars represent the comparison to the WT and the lines with represent direct comparisons between genotypes. n values reflect number of polytenes scored for each respective genotype. Chi-squared test, *** = p < 0.001.

Overall, it’s clear that the larger context of the histone gene array is not required for either CLAMP recruitment or specific function but that the local flanking context of the *H3/*H4p promoter is important for CLAMP function at the histone gene array.

Our results were somewhat surprising since other well-studied transcription factors rely exclusively on *cis* element sequence, rather than local context for function. For example, the *Drosophila* pioneer factor Zelda targets “TAGteam” sequences. Zelda competes with another TF, Grainyhead, for binding TAGteam *cis* elements and differences in the motif sequence elicits differential binding and function of Zelda (Harrison et al. 2010; Li and Eisen 2018). Because of this relationship between Zelda and specific *cis* element sequence, specific TAGteam sequences can be placed in combination in transgenes to titrate gene expression output (Li and Eisen 2018). Another example is the early *Drosophila* embryo factor Twist which binds a variety of *cis* element motifs. Twist targets several repetitive *cis* elements as well as E-box motifs, the sequences of which are not interchangeable: each E-box type corresponds to a discrete regulatory role (Ozdemir et al. 2011). However, unlike Zelda and Twist, CLAMP appears to glean significant information from the flanking local context wherein *cis* elements reside.

### CLAMP binding sequences from different loci do not functionally substitute in the context of the histone array in chromosomal females

The presence of CLAMP-recruiting elements within the local context of the *H3/H4*p is necessary for ectopic Mxc recruitment. Prior work demonstrated that the *H3/H4*p alone is sufficient to recruit HLB factors and even to initiate histone transcription, indicating that the rest of the array is dispensable at least for these actions (Salzler et al. 2013). However, when CLAMP is tethered to the *H3/H4*p in the absence of the GA-repeats, Mxc and other HLB factors are recruited, but transgenic histone transcription is not initiated (Rieder et al. 2017). Therefore, the specific CLAMP-GA-repeat interaction is critical for full CLAMP function at the histone genes. That being considered, CLAMP may integrate other local information that is unique to the large, repetitive endogenous histone locus. For example, CLAMP-binding sites exist in each of the ∼100 histone arrays (Bongartz and Schlossnig 2019), but it is unclear if CLAMP binds to all of these sites. Concentrating HLB factors at the locus likely contributes to important body properties such as phase separation (Hur et al. 2020) and facilitates histone biogenesis (Tatomer et al. 2016). The repetitive nature of the locus also likely leads to unique three-dimensional organization, both within and between loci (Carty et al. 2017; Fritz et al. 2018), which can impact the function of transcription factors such as CLAMP. Since these higher-order organizational aspects are not recapitulated at our transgenic histone gene arrays, we are unable to capture how they contribute to its contest-specific functions.

Given that CLAMP targets many GA-rich elements across the genome and that CLAMP may integrate local genomic context information to perform its functions, we next sought to investigate whether X-linked sequences can functionally substitute for the endogenous GA-repeats in the *H3/H4*p *in vitro*. We designed three hybrid DNA probes based on the sequence of the *H3/H4*p, but in which we replaced parts of the promoter with various amounts of X-linked sequence. In the “2 MRE” probe we replaced the two endogenous CLAMP recruiting GA-repeats with two 21 bp GA-rich MREs from the X-linked *roX2* gene. The “CES5C2” probe replaces the sequence between the *H3* and *H4* TATA boxes with sequence from the X-linked CES5C2 region containing three 21 bp GA-rich MREs. Finally, the “*roX2*” probe replaces the sequence between the *H3* and *H4* TATA boxes with sequence from the X-linked *roX2* CES containing four 21 bp GA-rich MREs (**Figure 4A**). We performed radiolabeled EMSAs using embryo extracts and all three hybrid probes robustly shifted (**Figure 4B**). CLAMP therefore binds X-linked GA-rich elements in the context of the *H3/H4*p *in vitro*, and this interaction appears similar to CLAMP binding of the endogenous GA-repeats.

**Figure 4:**
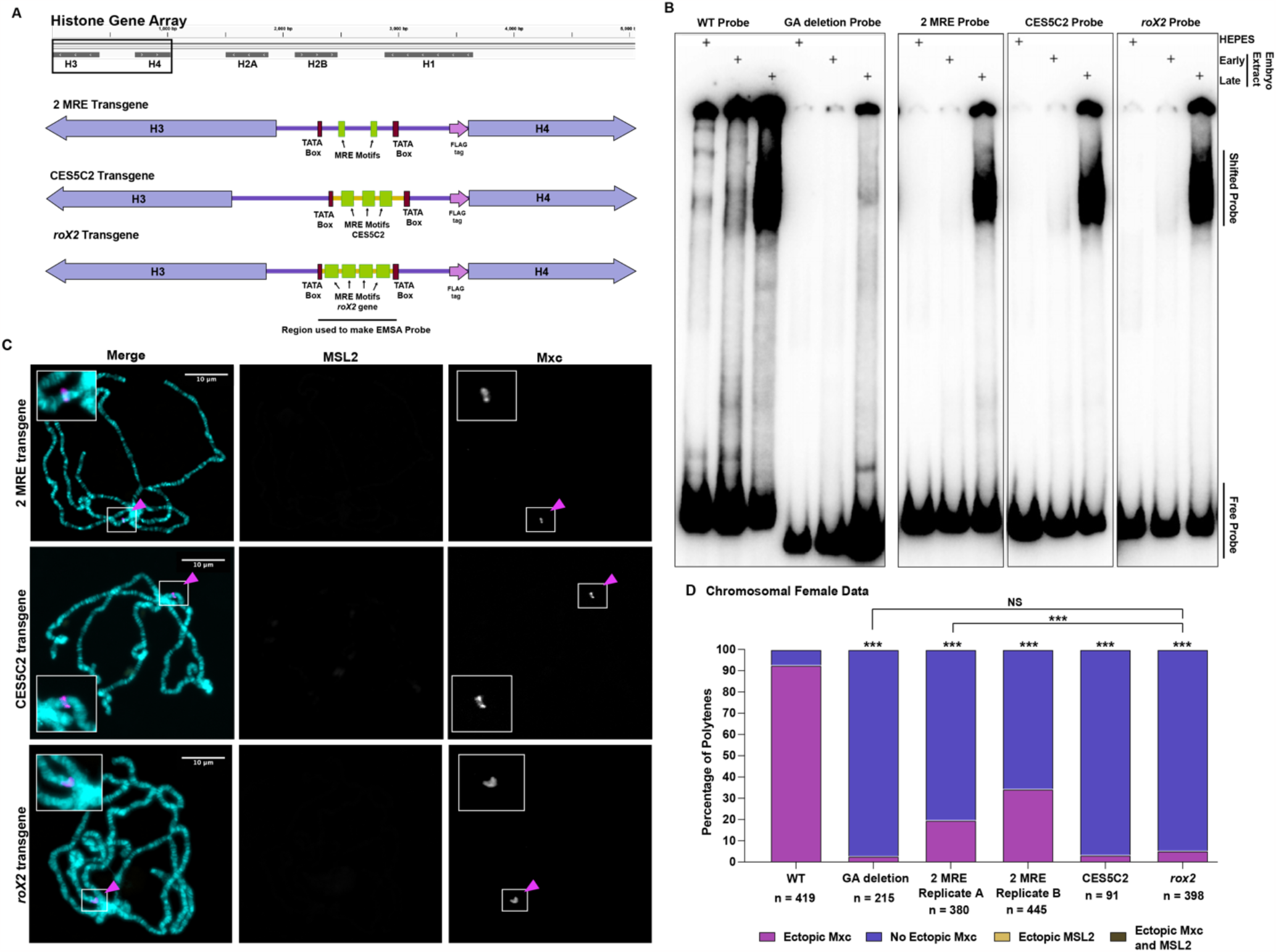
X-linked elements do not functionally substitute GA-repeats in the histone gene array in chromosomal females. **(A)** We engineered histone array transgenes in which we replaced parts of the *H3/H4p* sequence with varying amounts of X-linked sequence; the “2 MRE” transgenes replaces the GA-repeats with X-linked MREs from the *roX2* gene (light green boxes), the “CES5C2” transgene replaces the sequence between the TATA boxes (maroon) with the X-chromosome CES5C2 sequence (yellow line, light green bars indicate MREs), and the “*roX2”* transgene replaces the sequence between the TATA boxes with the CES sequence from the *roX2* gene (yellow line, light green bars indicate MREs). **(B)** We performed EMSA (gel shift) assays with early and late embryo extract (HEPES buffer serves as a control) and radiolabeled probes of the *H3/H4p* sequences of the histone array transgenes in (A) (WT: 226 bp, GA deletion: 198 bp, 2 MRE: 238 bp, CES5C2: 232 bp, *roX2*: 232 bp). Late embryo extract, containing CLAMP, shifts all EMSA probes of the *H3/H4*p other than the GA deletion probe. **(C)** We performed immunofluorescence staining on third instar larval polytene chromosomes from salivary glands in chromosomal females for Mxc (magenta) and MSL2 (yellow). DNA is stained with DAPI (cyan). Mxc localizes to the endogenous histone locus, which is used as an internal staining control (magenta arrow, solid outlined box). **(D)** Quantification of ectopic Mxc from polytene scoring shows a significant difference in the percentage chromosome spreads that have ectopic Mxc. Significance above the bars represent the comparison to the WT data and the lines with represent direct comparisons between datasets. n values reflect number of polytenes scored for each respective genotype. Chi-squared test, *** = p < 0.001

To translate our results *in vivo*, we engineered transgenic *Drosophila* lines carrying these hybrid transgenes with the same sequences as our EMSA probes (**Figure 4A**) and performed polytene chromosome immunostaining as above. We observed very little ectopic Mxc in all chromosomal female larvae regardless of which hybrid transgene they carried. Animals carrying the 2 MRE transgene were significantly less likely to show ectopic Mxc than the WT control animals, but significantly more likely to show ectopic Mxc compared to animals carrying the CES5C2 and *roX2* transgenes (**Figure 4C**,**D)**.

Together these data suggest that replacing just the *H3/H4*p GA-repeats impacts Mxc recruitment, in chromosomal females even though the majority of the *H3/H4*p sequence is preserved and CLAMP binds to this sequence *in vitro* (**Figure 4B**). Overall, our data from chromosomal females suggests that *cis* element sequence impacts CLAMP function at the transgenic histone gene array. Furthermore, in combination with our data showing the GA-repeats must reside in the *H3/H4*p for Mxc recruitment to the histone array transgene (**Figure 3)**, our results suggest that CLAMP is using cues from both *cis* element sequence and the local flanking regions of the *cis* element to determine its function at the transgenic histone gene array.

### X-linked sequences in the context of the histone gene array attract MSL2 in chromosomal males

Because *Drosophila* males undergo dosage compensation, MSLc members such as MSL2 are present in males whereas they are not expressed in females. When CLAMP binds the GA-rich MREs on the X-chromosome, it then recruits MSLc for dosage compensation (Soruco and Larschan 2014; Rieder et al. 2019). We therefore considered that our transgenes might behave differently in chromosomal males compared to females. We performed immunofluorescence staining on male polytene chromosomes with antibodies against Mxc and MSL2. Although MSL2 may have some DNA-binding capability (Villa et al. 2016), it requires CLAMP for efficient X-chromosome targeting (Soruco and Larschan 2014; Rieder et al. 2019).

Similar to our observations in chromosomal females, X-linked sequences do not functionally substitute in the local context of the *H3/H4*p in males: we observed few instances of ectopic Mxc in animals carrying the chimeric transgenes (**Figure 5B**,**C**). Strikingly, ectopic Mxc was significantly rarer in chromosomal males than in females carrying the 2MRE transgene (**Figure 5C**). However, in males we observed ectopic, autosomal MSL2: approximately 50% of polytene spreads from animals carrying the 2MRE transgene showed ectopic MSL2, whereas the majority of larvae carrying the CES5C2 or *roX2* transgene showed ectopic MSL2 recruitment (**Figure 5B**,**C)**. These results suggest that X-linked MRE or CES sequences recruit MSLc, even in the context of the transgenic histone gene array. These results confirm that CLAMP utilizes cues from the *cis* element sequence itself as well as the local flanking regions to determine its function at the transgenic histone gene array.

**Figure 5:**
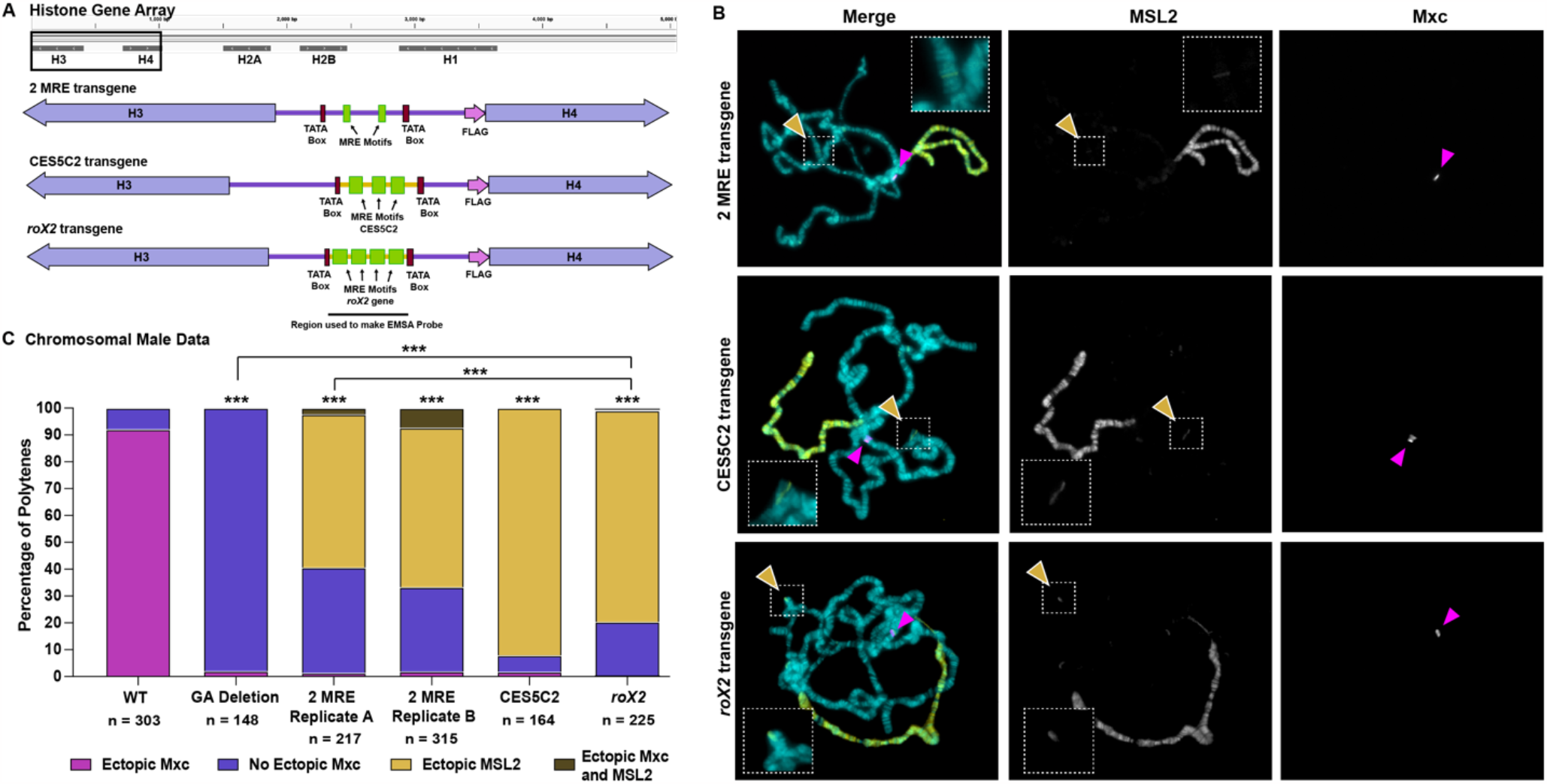
X-linked elements attract MSL2 in the context of the transgenic histone gene array in chromosomal males. **(A)** We engineered single copy histone array transgenes in which we replaced parts of the *H3/H4p* sequence with varying amounts of X-linked sequence as in Figure 4. **(B)** We performed immunofluorescence staining on third instar larval polytene chromosomes in chromosomal males for Mxc (magenta) and MSL2 (yellow). DNA is stained with DAPI (cyan). Mxc localizes to the endogenous histone locus, which is used as an internal staining control (magenta arrow). MSL2 marks the male X-chromosome and also serves as an internal staining control. Chromosomal males containing any of the three histone array transgenes with X-linked sequence show some amount of ectopic MSL2 staining on autosomes (white outlined yellow arrows, dotted white boxes). **(C)** Quantification of ectopic Mxc from polytene scoring shows a significant difference in the percentage chromosome spreads that have ectopic Mxc. Significance above the bars represent the comparison to the WT data and the lines with represent direct comparisons between genotypes. n values reflect number of polytenes scored for each respective genotype. Chi-squared test, *** = p < 0.001

When we replace the majority of the *H3/H4*p with X-linked sequence, CLAMP likely binds this region but performs its X-linked role, even in the context of the transgenic histone gene array, suggesting there may be other information within the local context, such as additional TF recruiting *cis* elements, that impact CLAMP histone locus role. This observation is interesting given that we recently discovered that in *Drosophila virilis*, a species ∼40 million years diverged from *D. melanogaster*, MSL2 is recruited to one of their two endogenous histone loci (Xie et al. 2022). The *D. virilis H3/H4*p carries poorly conserved, much shorter GA-repeats than those in *D. melanogaster* (Rieder et al. 2017), but CLAMP is still recruited to this sequence *in vitro* and, furthermore, recruits both Mxc and MSL2 to one of the two loci (Xie et al. 2022). This observation suggests that there may be evolutionary differences in CLAMP function and that there are different mechanisms for histone gene regulation, perhaps even within single species (Koreski et al. 2020). In addition, we recently explored other candidates that target the *D. melanogaster* histone gene array (Hodkinson et al. 2023). We identified several DNA-binding factors from this screen, including the Hox factor Ultrabithorax, that may provide CLAMP contextual cues for functioning at the histone locus. Ubx appears to interact specifically with the *H3/H4*p sequence and is therefore positioned close to the CLAMP binding sites. Given that there is likely a “secondary” mechanism to HLB formation that does not involve the CLAMP-GA-repeat interaction (Koreski et al. 2020); see below), other transcription factors emerge as a likely mechanism.

Overall, we show that the context-dependent transcription factor CLAMP incorporates both *cis* element sequence information as well as cues from local flanking context where its *cis* binding elements reside to govern its function. We show that CLAMP *cis* elements are not interchangeable and that, in the context of the histone locus, the local context of the *H3/H4*p provides CLAMP with critical cues to function at the histone genes. Together our findings provide new insights into our understanding of how TFs bind similar *cis* elements in locations across the genome but can preserve their specific regulatory functions at these each of these different loci.

## Acknowledgements

We would like to the members of the Rieder Lab as well as Dr. William Kelly and Dr. Roger Deal for their helpful remarks on the manuscript. We also would like to thank Dr. William Marzluf and Dr. Robert Duronio for Mxc antibody as well as Dr. Victoria Meller for MSL2 antibody. This work was supported by T32GM00008490 and F31HD105452 to LJH, K12GM00068 to CAS and HSC; F32GM140778 to CAS; and R00HD092625 and R35GM142724 to LER.

## Author contributions

Conceptualization: JG, CAS, and LER. Methodology: JG, LJH, CAS, PPD, TA, and LER. Validation: LJH, TA, and LER. Formal analysis: LJH, CAS, PPD, and LER. Investigation: LJH, CAS, and TA. Resources: TA and LER. Data curation: LJH and LER. Writing – Original Draft: LJH and LER. Writing – Review and Editing: LJH, JG, CAS, TA, and LER. Visualization: LJH and LER. Supervision: LER. Project Administration: LJH and LER. Funding Acquisition: LJH, CAS, and LER.

## Materials and methods

### Transgenics

Transgenes were constructed to include a 5 Kb histone array sequence consisting of the 5 replication-dependent histone genes and their relative promoters (McKay et al. 2015; Meers et al. 2018) in which the H4 and H2A genes are FLAG-tagged (24 bp) at the 5’ end to distinguish them from the endogenous histone genes. 500 bp DNA inserts containing the *H3/H4*p changes of interest were ordered from IDT and inserted via Gibson cloning (detailed above). All 1x histone array transgenes were inserted at the VK33 attP site on chromosome 3L (65B2) (Venken et al. 2006) using PhiC3-mediated integration (Groth et al. 2004) by GenetiVision (Houston, TX). Injected chimeric flies were crossed in pairs to a +/+; CyO/If ; TM3 (Sb) / TM6 (Tb) balancer stock. Resulting red-eyed progeny were selected and singly mated back to +/+; CyO/If ; TM3 (Sb) / TM6 (Tb) flies and a homozygous stock was established. Flies were maintained on standard cornmeal/molasses media at 18º C and transferred every 3-4 weeks.

### Electrophoretic Mobility Shift Assays

We performed EMSAs after Aoki et al. 2008 (Aoki et al. 2008) with minor modifications.

#### DNA probes

We made EMSA probes using PCR from gblocks (IDT) acquired during cloning as templates and the following primers: H3/H4 promoter F1: CACAGCACGAAAGTCACTAAAGAAC, H3/H4 promoter R1: GTTTGAAAACACAATAAACGATCAGAGC. We 5′ end labeled one pmol of probe with γ-32P-ATP (MP Biomedicals) using T4 polynucleotide kinase (New England BioLabs) in a 50 μl total reaction volume at 37°C for 1 hour. We used Sephadex G-50 fine gel (Amersham Biosciences) columns to separate free ATP from labeled probes. We adjusted the volume of the eluted sample to 100 μl using deionized water so that the final concentration of the probe was 10 fmol/μl.

#### Late embryo nuclear extracts

We prepared embryo extracts from 6-18hr Oregon R embryos collected on apple juice plates and aged 6 hours at room temperature. We performed nuclear extract preparation as in Aoki et al. 2008. We omitted the final dialysis step described in Aoki et al. and completed the extraction with the final concentration of KCl at 360 mM.

#### Shifts

We performed 20 μl binding reactions consisting of 0.5 μl (5 fmol) of labeled probe in the following buffer: 25 mM Tris-Cl (pH 7.4), 100 mM KCl, 1 mM EDTA, 0.1 mM dithiothreitol, 0.1 mM PMSF, 0.3 mg/ml bovine serum albumin, 10% glycerol, 0.25 mg/ml poly(dI-dC)/poly(dI-dC). We added 1 μl of nuclear extract and incubated samples at room temperature for 30 minutes. We loaded samples onto a 4% acrylamide (mono/bis, 29:1)-0.5× TBE-2.5% glycerol slab gel. We performed electrophoresis at 4°C, 180 V for 3-4 hours using 0.5× TBE-2.5% glycerol as a running buffer. We dried gels and imaged using a Typhoon 9410 scanner and Image Gauge software.

#### Supershifts

We preincubated reactions, including 5ug/ul poly(dA-dT)/poly(dA-dT), with OreR late nuclear extract (LNE) and antibodies for 30 min at room temperature before adding hot probe. We used 1 ul rabbit serum and 4 ul anti-CLAMP antibodies.

**Table 1:**
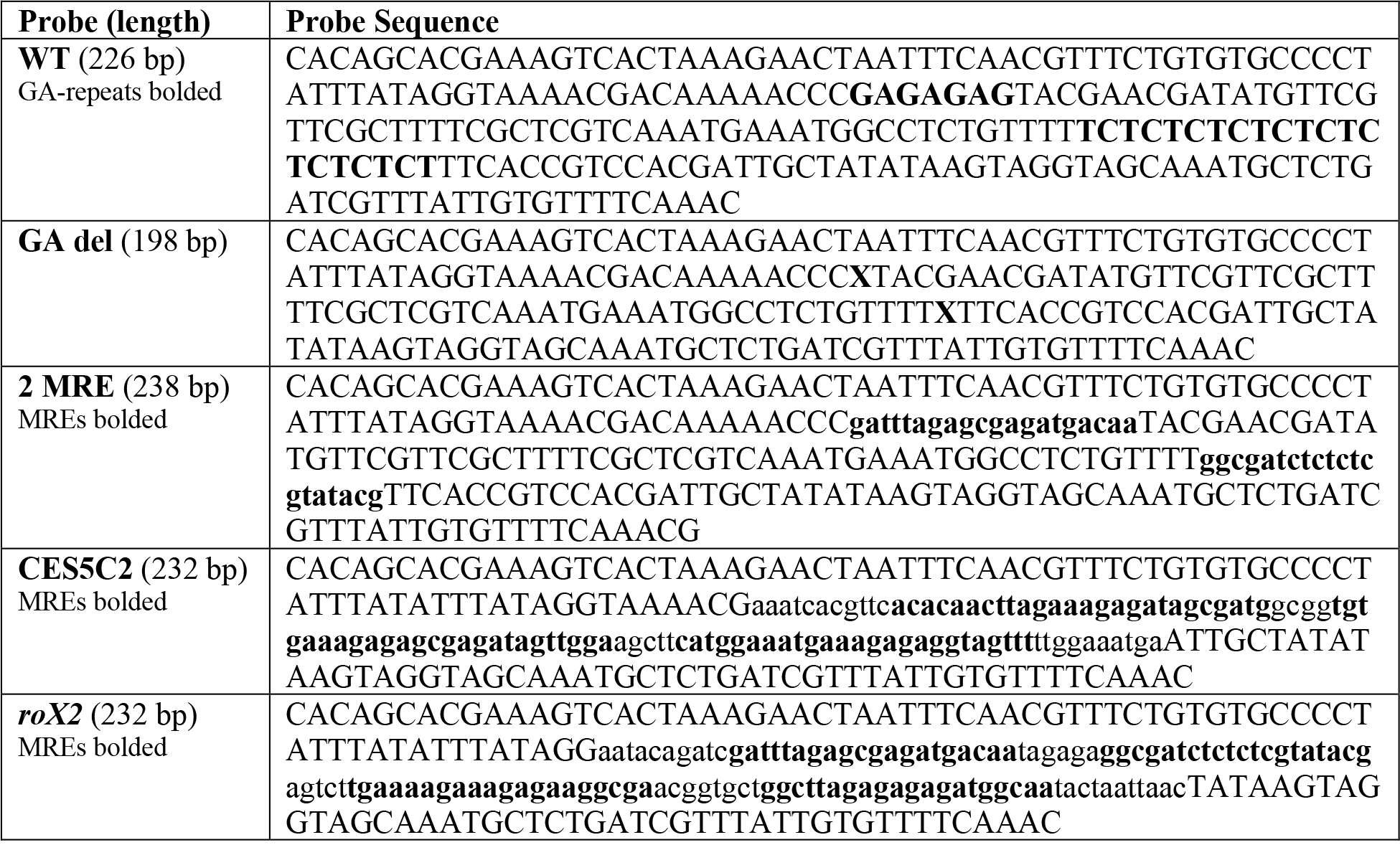
Sequences of EMSA probes.

### Polytene immunofluorescence and microscopy

We performed polytene chromosome squashes from salivary glands of sexed female and male third instar larvae. We passed glands through fix 1 (4% formaldehyde, 1% Triton X-100, in 1× PBS) for 1 min, fix 2 (4% formaldehyde, 50% glacial acetic acid) for 2 min, and 1:2:3 solution (ratio of lactic acid:water:glacial acetic acid) for 5 min prior to squashing and spreading. Slides were washed in 1X PBS, washed in 1% Triton X-100 (diluted in 1X PBS) and blocked for one hour in.5% BSA in 1X PBS. Slides were then incubated with primary antibody diluted in blocking solution (antibody specifics below) overnight at 4º C in a dark humid chamber. Slides were then washed in 1 X PBS and incubated with secondary antibody diluted in blocking solution (antibody specifics below) for two hours. Slides were then mounted using Prolong Diamond anti-fade reagent with DAPI (Thermo Fisher, P36961), and spreads were imaged on a using a Zeiss Scope.A1 equipped with a Zeiss AxioCam using a 40×/0.75 plan neofluar objective using AxioVision software. We used primary antibodies at the following concentrations: rabbit anti-CLAMP (1:1000; Novus/SDIX) (Larschan et al. 2012), guinea pig anti-Mxc (1:5000) (White et al. 2011, gift from the Duronio and Marzluff labs), and rabbit anti-MSL2 (1:150, a gift from the Meller Lab). We used AlexaFluor secondary 488 and 647 antibodies (Thermo Fisher Scientific) at a concentration of 1:1000.

### ChIP-seq data

CLAMP ChIP-seq was performed using protocol from Rieder et al. 2017 (Rieder et al. 2017) and data was taken from GEO (GSE152598, (Duan et al. 2021)). MSL3 ChIP-seq was performed using protocol from Rieder et al, 2017 and data was taken from GEO (GSE133637, (Rieder et al. 2019)). Mxc ChIP-seq was performed as in Rieder et al. 2017 using 2 ul guinea pig anti-Mxc antibody (gift from the Duronio/Marzluff laboratories).

### Data availability

Mxc ChIP-seq data is available on GEO (accession number pending).

### Bioinformatic analysis, alignment and visualization

We performed bioinformatics analysis as in Hodkinson *et al*. 2023 (Hodkinson et al. 2023). We directly imported individual FASTQ datasets into the web-based platform Galaxy (The Galaxy Community 2022) through the NCBI SRA Run Selector or using our sequencing files by selecting the desired runs and utilizing the computing Galaxy download feature. We retrieved the FASTQ files from SRA using the “Faster Download and Extract Reads in FASTQ format from NCBI SRA” Galaxy command. Because the ∼100 histone gene arrays are extremely similar in sequence (Bongartz and Schloissnig 2018), we do not utilize the dm6 or dm3 genomes and instead can collapse ChIP-seq data onto a single histone array (McKay et al. 2015; Bongartz and Schloissnig 2018; Koreski et al. 2020). We used a custom “genome” that includes a single Drosophila melanogaster histone array similar to that in Mckay et al. 2015, which we directly uploaded to Galaxy using the “upload data” feature, and normalized using the Galaxy command “NormalizeFasta” specifying an 80 bp line length for the output.fasta file. We aligned ChIP reads to the normalized histone gene array using Bowtie2 (Langmead and Salzberg 2012) to create .bam files using the user built-in index and “very sensitive end-to-end” parameter settings. We converted the .bam files to .bigwig files using the “bamCoverage” Galaxy command in which we set the bin size to 1 bp and set the effective genome size to user specified: 5000 bp (approximate size of l histone array). We also mapped relevant input or IgG datasets. If an input dataset was available, we normalized ChIP datasets to input using the “bamCompare” Galaxy command in which we set the bin size to 1 bp. We visualized the bigwig files using the Integrative Genome Viewer (IGV) (Robinson et al. 2011).

