## Supplemental Figure 1 for "Sequence reliance of a *Drosophila* context-dependent transcription factor"

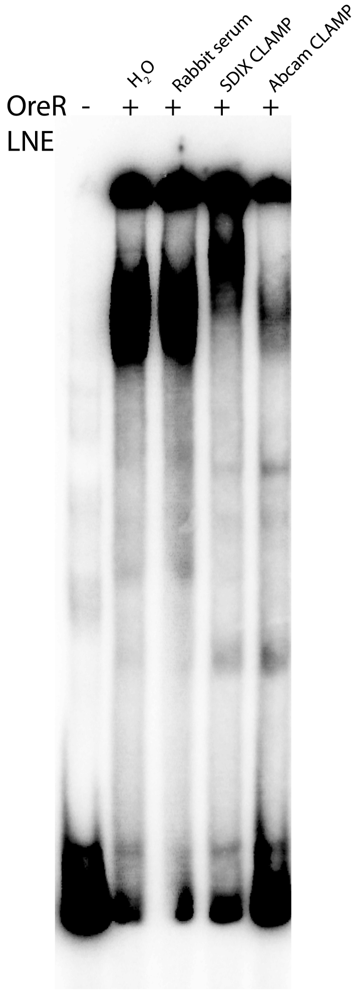


**Figure S1**: Supershift demonstrating that CLAMP is present in late nuclear extract (LNE) from wild type (OregonR) and shifts the *H3H4*p probe. Late nuclear extract shifts the *H3H4*p probe. Rabbit serum (negative control) does not supershift, while two anti-CLAMP antibodies from two different companies both supershift.
